# Zebrafish lack canonical necroptosis despite retaining functional RIPK3

**DOI:** 10.64898/2026.11.10.765635

**Authors:** Joana Furtado Milão, Lalita Ramakrishnan, Naoya Yamaguchi

## Abstract

Organismal development, homeostasis and immunity are facilitated by distinct forms of regulated cell death (RCD). These include conserved apoptosis and necroptosis, a proposed backup antimicrobial defence that requires RIPK3-mediated activation of MLKL and is predicted to be absent in several species lacking MLKL. Zebrafish studies have identified a RIPK3-dependent regulated necrosis that is distinct from necroptosis and is pathogenic in tuberculosis. Here, we experimentally confirm the bioinformatics-based prediction that zebrafish lack canonical necroptosis because they lack MLKL, despite retaining functional RIPK3. We take advantage of the finding that Caspase-8 promotes apoptosis while restraining necroptosis; its deficiency is embryonic lethal in mice unless RIPK3 or MLKL is also inhibited. In contrast, we found that Caspase-8-deficient zebrafish are viable through adulthood without obvious morphological defects. Inhibition of related caspases did not compromise viability, arguing against functional redundancy. Finally, Ripk3 overexpression failed to induce death in Caspase-8-deficient embryos. Our work highlights the zebrafish as a model for exploring RIPK3 and extrinsic apoptosis biology in disease contexts in the natural absence of canonical necroptotic execution.

**Summary statement:** Cell death drives many pathological conditions. Using *casp8* mutant zebrafish, we show that zebrafish lack canonical necroptosis despite retaining functional RIPK3.

## Introduction

Regulated cell death (RCD) is a genetically controlled process activated by diverse stimuli (Newton et al., 2024). Apoptosis, the first form of RCD to be characterized, is typically nonlytic and noninflammatory and contributes to organismal development and homeostasis throughout life (Newton et al., 2024). Its extrinsic pathway is initiated by death receptor activation and can be triggered by inflammatory stimuli (Newton et al., 2024, Tummers and Green, 2017). Other forms of RCD are lytic, causing necrotic cell death, and can both be triggered by and promote inflammation (Orning and Lien, 2021). These pathways can support immunity by eliminating infected cells and limiting pathogen expansion (Kaiser et al., 2013). Both bacterial and viral pathogens have evolved mechanisms to inhibit RCD (Faherty and Maurelli, 2008, Kaiser et al., 2013). Some inhibit Caspase-8, which initiates extrinsic apoptosis downstream of death receptors (Fig. 1A)(Kaiser et al., 2013). Necroptosis is an alternative lytic form of RCD that is *inhibited* rather than initiated by Caspase-8 (Orning and Lien, 2021, Tummers and Green, 2017) (Fig. 1A). This arrangement can allow infected cells to undergo necroptosis when pathogens inhibit Caspase-8; accordingly, necroptosis is proposed to have evolved as a backup against pathogen-mediated inhibition of apoptosis (Kaiser et al., 2013). The importance of Caspase-8 in restraining necroptosis is illustrated by the embryonic lethality of Caspase-8-deficient mice, which is rescued by loss of either the kinase RIPK3 or its downstream effector, the pseudokinase MLKL (Fig. 1A)(Oberst et al., 2011, Kaiser et al., 2011, Alvarez-Diaz et al., 2016). In humans, pathogenic biallelic CASP8 variants cause immunodeficiency and immune dysregulation, including lymphoproliferation and intestinal inflammation (Chun et al., 2002). Despite the role ascribed to necroptosis in host defence, RIPK3 and MLKL have each been lost repeatedly during vertebrate evolution, whereas RIPK1, a regulator of cell survival and extrinsic apoptosis, is widely conserved (Fay et al., 2025, Dondelinger et al., 2016).

**Figure 1:**
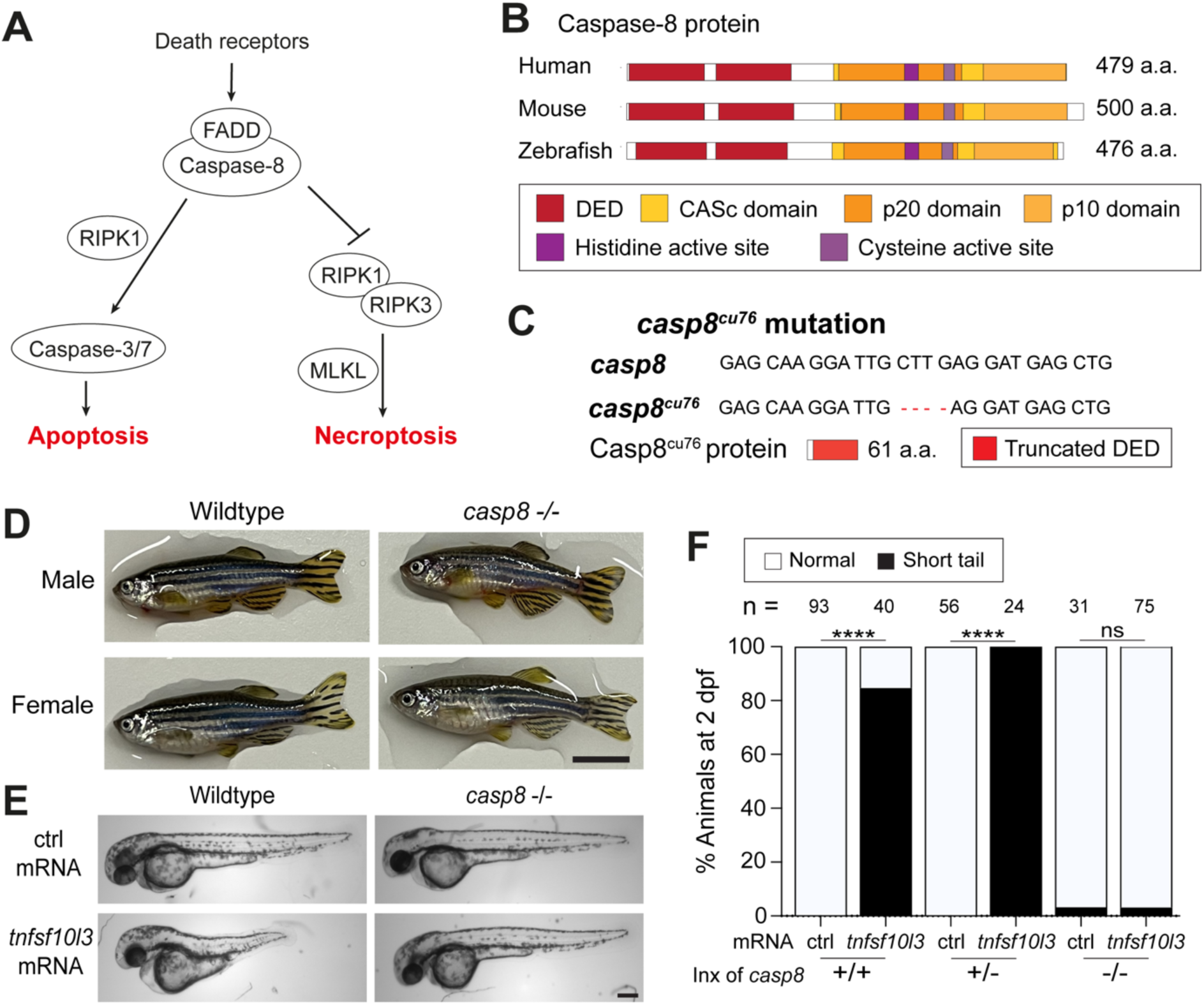
Generation and characterization of *casp8* mutant zebrafish. (A) Diagram illustrating Caspase-8-mediated regulation of apoptosis and necroptosis. (B) Schematic representation and domain architecture of human, mouse and zebrafish Caspase-8 proteins. (C) *casp8^cu76^*mutant allele and its protein product. (D) Wild-type and *casp8^cu76^*homozygous adults. Scale bar: 1 cm. (E) Larvae with Tnfsf10l3-mediated short tail phenotype and its rescue in maternal-zygotic *casp8^cu76^* homozygous background. Scale bar: 250 μm. (F) Scores for larvae with Tnfsf10l3-mediated short tail phenotype at 2 dpf derived from indicated crosses. Inx: intercross. ctrl: *emiRFP670* mRNA. ***: *p*<0.0001 and ns: *p*>0.05 with Fisher’s exact test.

Zebrafish (*Danio rerio*) offer genetic, pharmacological and imaging tools for studying RCD in a whole-animal vertebrate model of human disease (Patton and Tobin, 2019, Pagan et al., 2022). Many components of the RCD machinery are conserved in zebrafish (Inohara and Nunez, 2000, Krumschnabel and Podrabsky, 2009). However, zebrafish lack an MLKL orthologue while retaining RIPK3, which has been implicated in cell death in several disease contexts (Roca and Ramakrishnan, 2013, Jim et al., 2022, Yang et al., 2020, Roca et al., 2019, Viringipurampeer et al., 2014). We previously showed that TNF excess drives RIPK3-dependent pathogenic necrosis of mycobacterium-infected zebrafish and human macrophages through a pathway distinct from canonical necroptosis (Roca and Ramakrishnan, 2013, Roca et al., 2022, Roca et al., 2019). These observations raise the question of whether zebrafish additionally retain a RIPK3-dependent, Caspase-8-regulated necroptotic pathway using an unrecognized MLKL homologue or an alternative effector.

Here, we show that zebrafish completely lacking Caspase-8 survive to adulthood and that RIPK3 overexpression fails to induce Caspase-8-independent embryonic death. Our findings support the predicted absence of canonical necroptosis in zebrafish and establish a model for studying extrinsic apoptosis and RIPK3 functions outside this pathway.

## Results

### Caspase-8 is required for extrinsic apoptosis but dispensable for zebrafish viability

Caspase-8 promotes extrinsic apoptosis while restraining necroptosis, and in mice, its deficiency is embryonic lethal owing to uncontrolled necroptosis (Sakamaki et al., 2002, Varfolomeev et al., 1998). We reasoned that if canonical necroptosis is operant in zebrafish then Caspase-8 loss should cause early lethality. To test this, we used CRISPR-Cas9 to generate the zebrafish *casp8^cu76^* (previously named *casp8a*, Fig. 1B) allele, which harbors a 4 bp deletion in exon 2, creating a frameshift and a premature stop codon within the first death effector domain (DED) (Fig. 1C). Homozygous *casp8^cu76^* mutants were recovered at Mendelian ratios and survived to adulthood without obvious morphological defects (Fig. 1D; Fig. S1A). We did not detect nonsense-mediated decay of mutant *casp8* mRNA (Fig. S1B), arguing against genetic compensation triggered by mutant mRNA decay (El-Brolosy et al., 2019, Rossi et al., 2015, Ma et al., 2019) as an explanation for viability.

Zebrafish undergo Caspase-8-dependent extrinsic apoptosis (Eimon et al., 2006). Injection *tnfsf10l3* (previously called *DL1b* and *tnfsf10-3*) mRNA into one-cell-stage embryos causes notochord cell apoptosis and a short-tail phenotype that can be scored at 2 days post-fertilization (dpf) (Eimon et al., 2006). To test whether *casp8^cu76^* abolished Caspase-8 function, we injected *tnfsf10l3* mRNA into embryos from wild-type crosses or *casp8^cu76^* heterozygous intercrosses. All injected progeny developed the short-tail phenotype regardless of genotype, suggesting that Caspase-8 activity persisted in the mutants (Fig. 1F). This could reflect maternally supplied *casp8* mRNA and/or Caspase-8 protein (Sakata et al., 2007). We therefore injected *tnfsf10l3* mRNA into embryos from *casp8^cu76^* homozygous intercrosses. These maternal-zygotic mutants were completely protected from the short-tail phenotype, confirming that *casp8^cu76^* is a loss-of-function allele (Fig. 1E, F). Together, these findings establish that Caspase-8 is required for extrinsic apoptosis but dispensable for zebrafish viability. Mutant survival could reflect inhibition of necroptosis by redundant factors or an absence of canonical necroptosis.

### Caspase-10 disruption and CrmA expression do not compromise zebrafish viability

Caspase-10 was our leading candidate for compensating for Caspase-8 loss. Although absent in mice, it is present in humans (Sakamaki et al., 2015), where it can act as an initiator caspase in extrinsic apoptosis and shares several substrates with Caspase-8 (Fig. S2A)(Fischer et al., 2006, Chang et al., 2002, Kischkel et al., 2001, Milhas et al., 2005, Takahashi et al., 2006). We first tested whether zebrafish Caspase-10 could induce cell death. Injection of *casp10* mRNA into one-cell-stage embryos induced death to a similar extent as *casp8* mRNA (Fig. 2A). Next, we used G0 CRISPR-Cas9 mutagenesis (Wu et al., 2018) to generate somatic *casp10* mutations in wild-type or maternal-zygotic *casp8* mutant embryos and asked whether this compromised viability. Larvae from both backgrounds showed similar survival over five days and no overt morphological defects (Fig. 2B, C). These results argue against Caspase-10 compensating for Caspase-8 loss to maintain larval viability.

**Figure 2:**
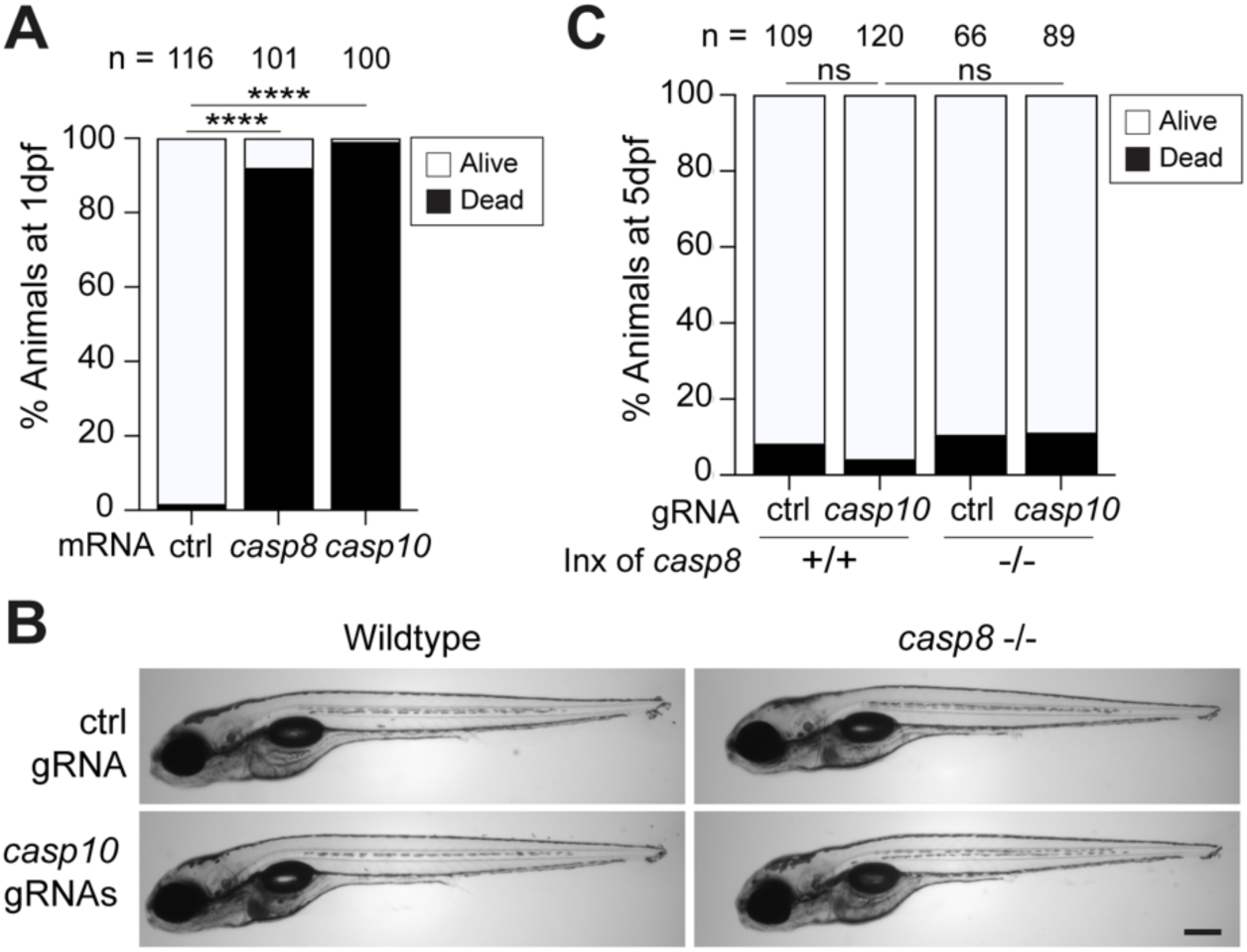
Survival of *casp8; casp10* G0 CRISPR double mutant larvae. (A) Percentage of dead embryos at 1 dpf injected with indicated mRNA. ctrl: *emiRFP670* mRNA. ****: *p*<0.0001 with Fisher’s exact test. (B) *casp10* or *aicda* (control) Cas9 RNP-injected wild-type or *casp8* mutant larva at 5 dpf. Scale bar: 250 μm. (C) Percentage of dead larvae at 5 dpf in wild-type or *casp8* mutant background injected with *casp10* or *aicda* (control) Cas9 RNPs. Inx: intercross. ns: *p*>0.05 with Fisher’s exact test.

Genome duplication in zebrafish (Howe et al., 2013) raised the possibility that additional caspase-8-like genes could compensate for the loss of *casp8* and *casp10*. For example, zebrafish *casp8l1* (previously named *casp8b* and *casp-8a-like*) encodes a protein with C-terminal similarity to Caspase-8 and Caspase-10 but lacking DED motifs (Spead et al., 2018) (Fig. S2A). Injection of *casp8l1* mRNA into wild-type embryos also induced death, demonstrating pro-death activity similar to that of Caspase-8 and Caspase-10 (Fig. 3A).

**Figure 3:**
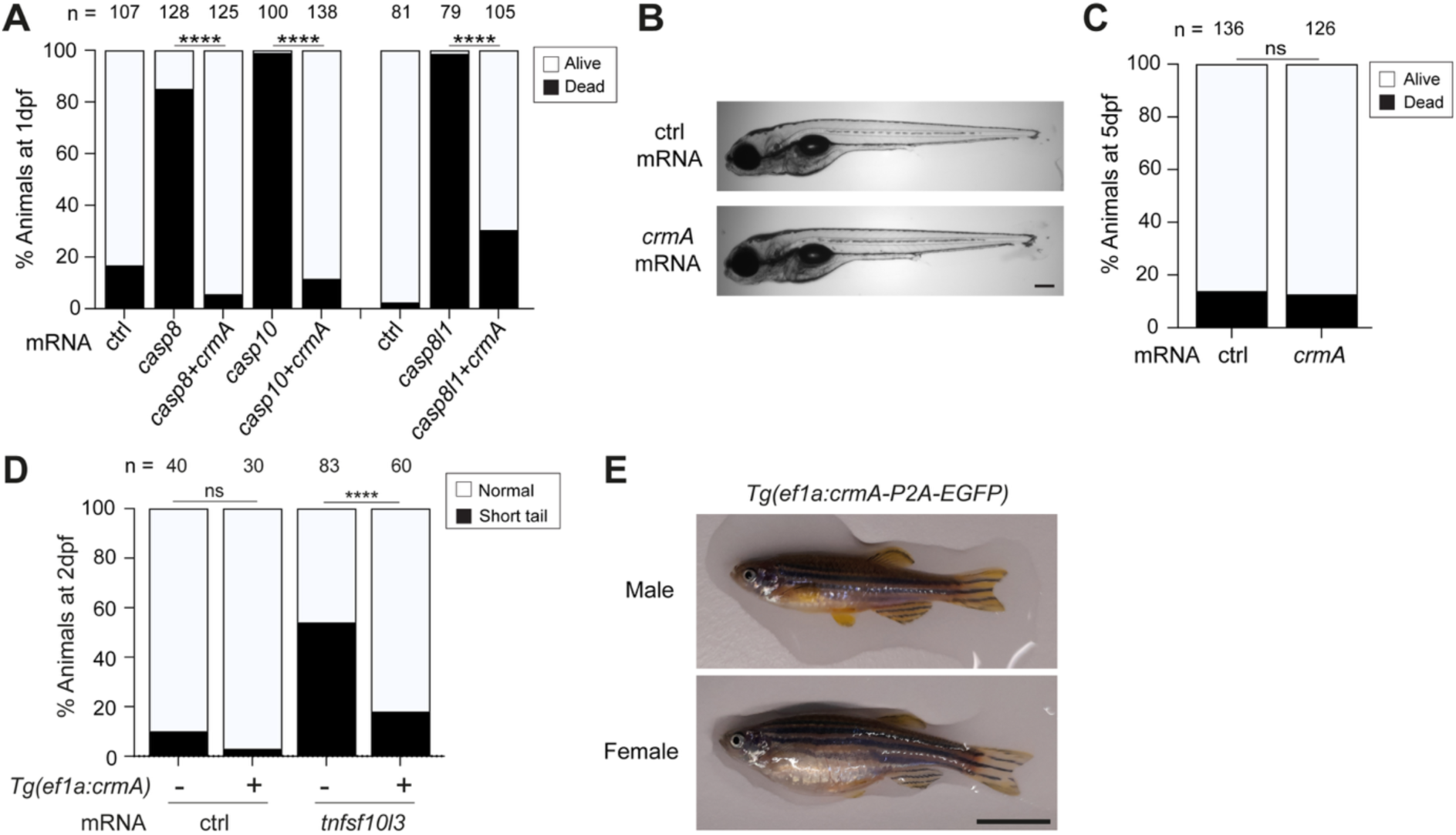
Survival of larvae with simultaneous inhibition of caspase-8-like proteins by CrmA. (A) Percentage of embryos dead at 1 dpf injected with mRNA from *casp8*-like genes with or without *crmA* mRNA. ctrl: *emiRFP670* mRNA. ****: *p*<0.0001 with Fisher’s exact test. (B) Larva injected with *crmA* or *emiRFP670* (control) mRNA at 5 dpf. Scale bar: 250 μm. (C) Percentage of larvae dead at 5 dpf injected with *crmA* or *emiRFP670* (control) mRNA. ns: *p*>0.05 with Fisher’s exact test. (D) Scores for larvae with Tnfsf10l3-mediated short tail phenotype at 2 dpf with or without CrmA expressed from the transgene. ctrl: *emiRFP670* mRNA. ****: *p*<0.0001 and ns: *p*>0.05 with Fisher’s exact test. (E) *Tg(ef1a:crmA-P2A-EGFP)^cu78^* transgenic adults. Scale bar: 1 cm.

Testing whether Caspase-8l1 compensates for Caspase-8 and Caspase-10 would require generating a *casp8*;*casp10*;*casp8l1* triple mutant and would not exclude redundancy involving other genes. To inhibit several related caspases simultaneously, we used cytokine response modifier A (CrmA), a potent Caspase-8 inhibitor from cowpox virus (Garcia-Calvo et al., 1998, Zhou et al., 1997). In susceptible mammalian cells, CrmA can shift cell death from extrinsic apoptosis to necroptosis (Holler et al., 2000, Ma et al., 2005, Vercammen et al., 1998, Oberst et al., 2011). We confirmed that CrmA suppressed death induced by Caspase-8, Caspase-10 and Caspase-8l1 by co-injecting *crmA* with the respective caspase mRNAs into one-cell-stage embryos (Fig. 3A). This established that CrmA inhibited the pro-death activity of all three zebrafish caspases. We then injected *crmA* mRNA alone and observed neither excess death nor morphological abnormalities (Fig. 3B, C) (Eimon et al., 2006). Because injected mRNA can be short-lived, we tested sustained inhibition by generating *Tg(ef1a:crmA-P2A-EGFP)^cu78^* transgenic zebrafish that ubiquitously expressed CrmA and EGFP (Fig. S3). Transgenic CrmA suppressed the Tnfsf10l3-induced short-tail phenotype, confirming functional inhibition of Caspase-8-dependent apoptosis (Fig. 3D). The transgenic animals survived to adulthood and appeared normal (Fig. 3E). Together, these results argue against redundancy among the tested caspases as the explanation for *casp8* mutant viability and support the absence of canonical necroptosis.

### Somatic *fadd* disruption does not cause early lethality

To corroborate these findings, we tested the effect of *fadd* disruption in zebrafish. FADD is an adaptor protein that links death receptors to Caspase-8 and Caspase-10 (Vincenz and Dixit, 1997, Fu et al., 2016) (Fig. 1A). It is required for both Caspase-8-mediated extrinsic apoptosis and Caspase-8-dependent inhibition of necroptosis (Zhang et al., 2011). Accordingly, *Fadd*-deficient mice, like Caspase-8-deficient mice, die prenatally (Zhang et al., 1998, Yeh et al., 1998). Zebrafish treated with a *fadd*-targeting morpholino have been reported to survive normally (Eimon et al., 2006). To extend this observation using an independent approach, we generated somatic *fadd* mutations by G0 CRISPR-Cas9 mutagenesis. At 5 dpf, 96% of *fadd*-targeted G0 larvae survived without overt morphological defects (Fig. 4A, B). This result provides independent support for the absence of a Fadd- and Caspase-8-regulated necroptotic death pathway during zebrafish development.

**Figure 4:**
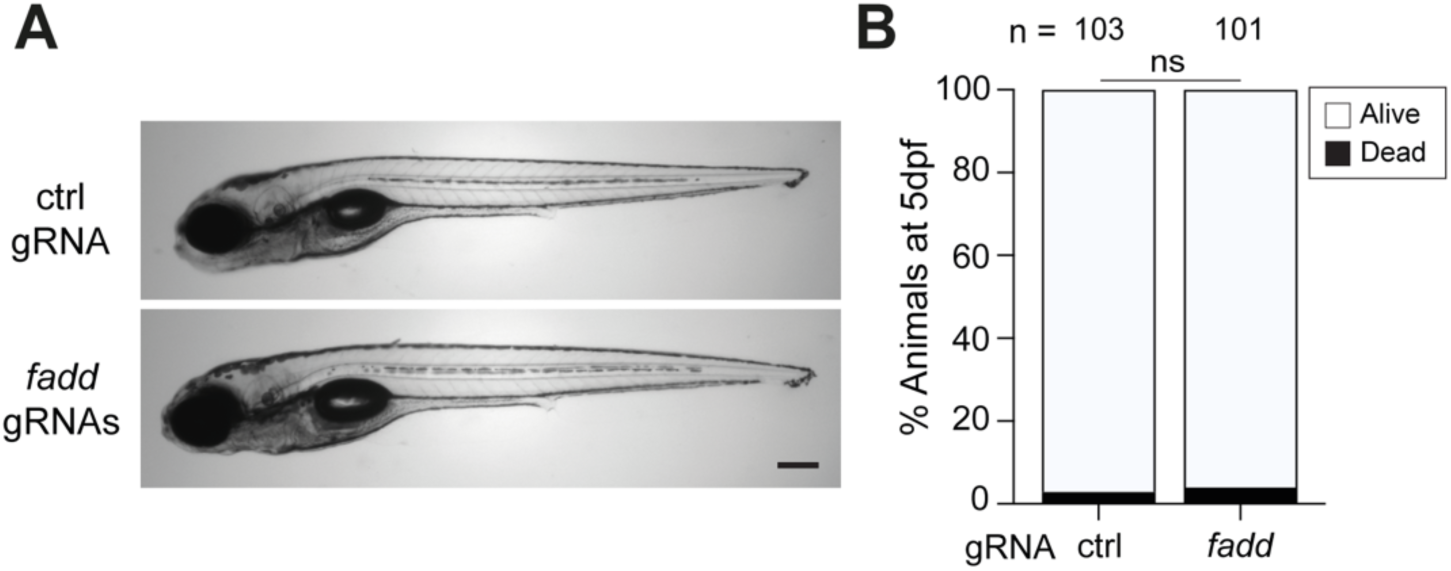
Survival of *fadd* G0 CRISPR mutant larvae. (A) *fadd* or *aicda* (control) Cas9 RNP-injected wild-type larva at 5 dpf. Scale bar: 250 μm. (B) Percentage of dead larvae at 5 dpf injected with *fadd* or *aicda* (control) Cas9 RNPs. ns: *p*>0.05 with Fisher’s exact test.

### Ripk3 overexpression does not kill embryos whereas Ripk1l induces Caspase-8-dependent death

RIPK3 can mediate necrosis independently of MLKL, and the resulting cell death can be difficult to distinguish morphologically from necroptosis (Zhang et al., 2016). We previously identified a RIPK3-dependent necrotic pathway that selectively affects mycobacterium-infected macrophages and is distinct from canonical RIPK3-MLKL-dependent necroptosis (Roca and Ramakrishnan, 2013, Roca et al., 2022, Roca et al., 2019). In a zebrafish model of insulin resistance, overnutrition causes β-cell loss involving ER stress, Ripk3-dependent *il1b* induction and macrophage recruitment; similar inflammatory responses were observed in mouse β-cells and human islets (Yang et al., 2020). RIPK3-dependent cone photoreceptor death has also been reported in *pde6c* mutant zebrafish, although its downstream execution mechanism remains unresolved (Viringipurampeer et al., 2014).

The viability and normal development of Caspase-8-deficient zebrafish supported the absence of canonical necroptosis. As a complementary approach, we asked whether excess Ripk3 could induce Caspase-8-independent embryonic death, including a possible MLKL-independent form of regulated necrosis. Injection of *ripk3* mRNA at the one-cell stage did not cause embryonic lethality (Fig. 5A). Injection of *ripk3*-sfGFP mRNA confirmed protein production yet likewise caused no death (Fig. S4A, B). Because interaction between RIPK1 and RIPK3 can activate RIPK3 during necroptotic signaling (Li et al., 2012, Cho et al., 2009), we next co-injected *ripk1l*, encoding the zebrafish RIPK1 ortholog, and *ripk3* mRNAs. Co-overexpression caused embryonic lethality, raising the possibility of a RIPK1-RIPK3-dependent death pathway. If this death reflected necroptosis normally restrained by Caspase-8, loss of Caspase-8 would not be expected to rescue it. Instead, lethality was rescued by using maternal-zygotic *casp8*^-/-^ embryos (Fig. 5A). These results argued against necroptosis as the cause of the observed death and were consistent with Caspase-8-dependent apoptosis (Peterson et al., 2017, Zhang et al., 2019). To determine whether Ripk3 co-overexpression was required, we injected *ripk1l* mRNA alone. This caused similar lethality to *ripk1l* and *ripk3* co-injection, and death was rescued by co-injection of *crmA* mRNA (Fig. 5B) or loss of Caspase-8 (Fig. 5C). Thus, Ripk1l overexpression induced Caspase-8-dependent death without requiring Ripk3 co-overexpression. Together, these experiments provide further evidence against a functional canonical necroptotic pathway in zebrafish.

**Figure 5:**
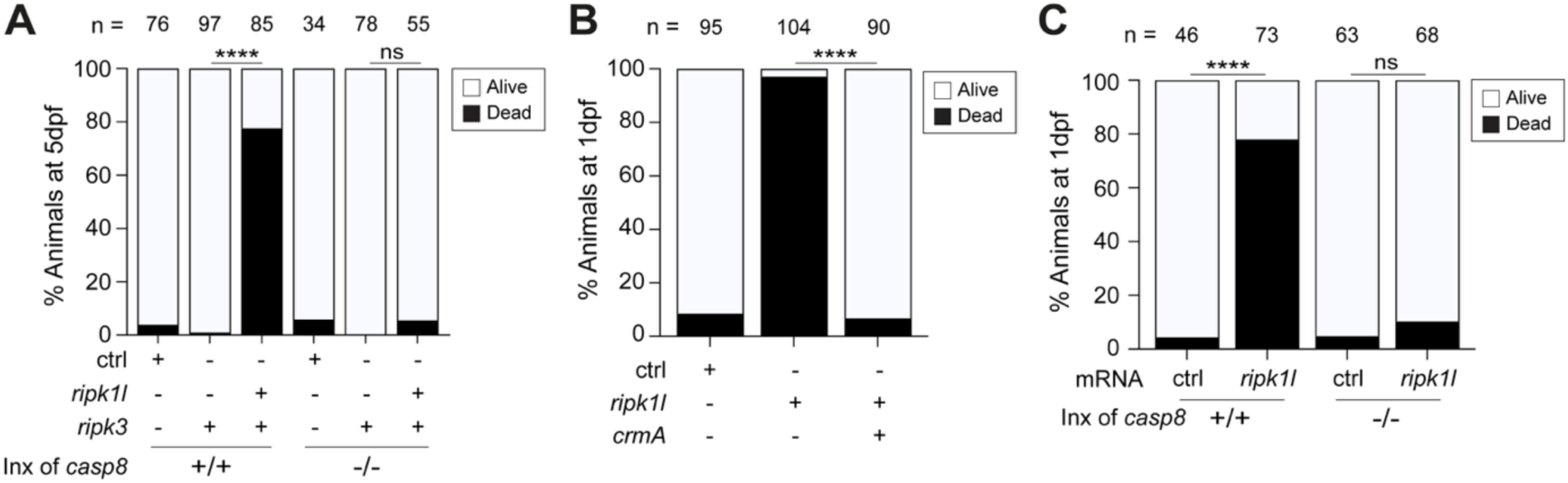
Caspase-8-dependent death caused by Ripk1l, but not by Ripk3. (A) Percentage of larvae dead at 5 dpf from indicated crosses injected with combinations of *ripk1l*, *ripk3* and ctrl mRNAs. Inx: intercross. ctrl: *emiRFP670* mRNA. ***: *p*<0.0001 and ns: *p*>0.05 with Fisher’s exact test. (B) Percentage of embryos dead at 1 dpf injected with *ripk1l* mRNA with or without *crmA* mRNA. ctrl: *emiRFP670* mRNA. ****: *p*<0.0001 with Fisher’s exact test. (C) Percentage of embryos from indicated crosses and dead at 1 dpf by injection of *ripk1l* or ctrl mRNA. Inx: intercross. ctrl: *emiRFP670* mRNA. ****: *p*<0.0001 and ns: *p*>0.05 with Fisher’s exact test.

## Discussion

Our multiple independent approaches support the bioinformatically predicted absence of canonical necroptosis in zebrafish, which lack MLKL. This finding is important given the increasing use of the zebrafish for studies of RCD in human disease modeling (White and Patton, 2023, Patton and Tobin, 2019).

RIPK1, which regulates cell survival and death, is widely conserved, whereas RIPK3 and MLKL have each been repeatedly lost during vertebrate evolution with an apparently patchy distribution. MLKL is absent from *Carnivora* but retained in several fish species (Dondelinger et al., 2016, Fay et al., 2025). These patterns could reflect species-specific pathogen pressures or compensation by other RCD pathways (Dondelinger et al., 2016). Known RIPK3-dependent responses in zebrafish, including islet inflammation and TNF-driven necrosis of mycobacterium-infected macrophages, have counterparts in human cells (Roca et al., 2019, Yang et al., 2020). These responses are therefore not zebrafish-specific innovations, although compensatory reliance on existing pathways remains possible.

Our findings show that loss of Caspase-8-dependent extrinsic apoptosis is compatible with adult viability. This is consistent with the survival of *Casp8*-deficient mice when *Ripk3* or *Mlkl* is also removed, preventing unrestrained necroptosis (Oberst et al., 2011, Kaiser et al., 2011, Alvarez-Diaz et al., 2016). Mice expressing Caspase-8(D387A), which impairs extrinsic apoptosis while retaining necroptosis suppression, also survive to adulthood (Oberst et al., 2011, Kang et al., 2008).

Caspase-8 loss affects nonapoptotic functions as well as extrinsic apoptosis, and additional mutations can alter RCD pathway interactions (Orning and Lien, 2021). Zebrafish permit analysis of Caspase-8 deficiency without further genetic disruption of the necroptotic machinery. They therefore provide a complementary whole-animal model for dissecting Caspase-8-dependent apoptosis and RIPK3 biology outside canonical necroptosis.

## Materials and methods

### Ethics

Zebrafish husbandry and experiments were conducted in compliance with guidelines from the UK Home Office using protocols approved by the Animal Welfare and Ethical Review Body of University of Cambridge.

### Zebrafish husbandry

Zebrafish embryos and larvae of the wild-type AB strain (Zebrafish International Resource Center: ZIRC) were used in experiments. Embryos were staged as previously described (Kimmel et al., 1995). Embryos were incubated at 28.5°C from one-cell stage (0 hours-post-fertilization: hpf) until the indicated time. All experiments were conducted in larvae such that their sex was undetermined. The general conditions for maintaining adult zebrafish were described before (Pagan et al., 2022). Zebrafish embryos were collected and kept in egg water (0.18 g/L Instant Ocean Salt and 0.25 μg/mL methylene blue) at 28.5 °C without the day-night cycle.

### Generation of the *casp8^cu76^* mutant by CRISPR-Cas9

The *casp8^cu76^* mutant was generated by CRISPR-Cas9 gene editing following the method described before (Hwang et al., 2013). Briefly, the DR274 plasmid (Addgene #42250) was initially digested with BsaI restriction enzyme (New England Biolabs, Hitchin, U.K.) and cloned with annealed nucleotides harboring a *casp8*-specific targeting sequence (F: 5’-TAGGATGAGCAAGGATTGCTTG-3’, R: 5’-AAACCAAGCAATCCTTGCTCAT-3’). Subsequently, this *DR274-casp8* sgRNA vector was linearized with DraI restriction enzyme (New England Biolabs) and used as a template for *in vitro* transcription using T7 RNA Polymerase (New England Biolabs). The transcribed gRNA was purified with RNeasy Mini Kit (Qiagen, Hilden, Germany) and eluted in RNase Free Water. *cas9* mRNA was prepared by the *in vitro* transcription from the linearized *pCS2-nCas9* (Addgene, MA, U.S.A., #47929) (Jao et al., 2013) using the mMESSAGE mMACHINE SP6 transcription Kit (Thermo Fisher Scientific, MA, U.S.A.). 1-2 nL of the injection mix containing 12.5 ng/μL of *casp8* gRNA, 300 ng/μL of *cas9* mRNA and 0.5% Phenol Red was injected into one-cell-stage embryos. CRISPR-injected embryos were raised to adulthood and subsequently out-crossed to wild-type fish to screen germline mutations. A *casp8* mutant allele harboring a 4-base pair (bp) deletion in the exon 2 was selected as *casp8^cu76^* for the subsequent studies. Genotyping of *casp8^cu76^* was performed using Kompetitive Allele Specific PCR (KASP)-based assay (LGC Biosearch Technologies, Middlesex, U.K.) in a CFX Connect thermocycler (Bio-Rad, CA, U.S.A.) using the primers listed in Table S1.

The mutation was verified by reverse transcribing the whole coding sequence of *casp8^cu76^* from its mRNA. mRNAs from F3 generation of *casp8^cu76^* embryos were harvested using RNeasy Mini Kit (Qiagen) and cDNA was synthesized using PrimeScript 1^st^ Strand cDNA Synthesis Kit (Takara Bio, Shiga, Japan). Subsequently, *casp8* coding sequence was amplified using primers listed in Table S1 and cloned into *pCS2(+)* plasmid using Gibson assembly followed by whole plasmid sequencing.

### Quantitative RT-PCR

Nonsense-mediated decay of *casp8*^cu76^ mRNA was tested by quantitative RT-PCR (qRT-PCR). Briefly, a part of the fin from 4 day post-fertilization (dpf) embryos derived from *casp8*^cu76^ heterozygous intercross was dissected out and genomic DNA was extracted from the dissected fin sample using the HotShot method. Using genomic DNA, the genotype of individual larva was determined, and 6 larvae were pooled based on their genotype. RNA was extracted from pooled larval samples using RNeasy Mini Kit (Qiagen) and cDNA was prepared by PrimeScript 1^st^ Strand cDNA Synthesis Kit (Takara Bio). qRT-PCR was performed in a CFX Connect thermocycler (Bio-Rad) using the primers targeting *casp8* and *actb2* as a control. Used primers are listed in Table S1. *casp8* mRNA abundance was quantified using the ΔΔCt method normalized by *actb2*.

### Plasmid preparation

The *pEF6-crmA* plasmid was purchased from NovoPro Bioscience Inc. (Shanghai, China). The coding sequences of the zebrafish genes *casp8*, *casp10*, *casp8l1*, *ripk1l,* and *tnfsf10l3* were cloned into *pCS2(+)* backbone with Gibson assembly using PCR primers listed in Table S1 and zebrafish cDNA prepared from 1 dpf AB embryos using PrimeScript 1^st^ Strand cDNA Synthesis Kit to generate the plasmids *pCS2(+)-casp8*, *pCS2(+)-casp10*, *pCS2(+)-casp8l1*, *pCS2(+)-ripk1l*, and *pCS2(+)-tnfsf10l3*. *pCS2(+)-ripk3* and *pCS2(+)-ripk3-sfGFP* were cloned using the coding sequence of zebrafish *ripk3* obtained from IDT as a gBlock fragment and the coding sequence of *super-folder GFP* (*sfGFP*) with Gibson assembly. Similarly, *pCS2(+)-emiRFP670* was generated to prepare control mRNA from *pH2B-emiRFP670* (Addgene#136571) (Matlashov et al., 2020). *pTol2-ef1a-crmA-P2A-EGFP* was generated by Gibson assembly using *crmA* coding sequences amplified from the *pEF6-crmA* plasmid and the *pTol2-ef1a-EGFP* backbone vector amplified from the *pTol2-ef1a-MLS-EGFP* plasmid (Addgene#31241) (Kim et al., 2008). All plasmids generated were verified by whole plasmid sequencing.

### *In vitro* transcription and microinjection of mRNAs

DNA templates for *in vitro* mRNA transcription were generated by PCR using primers listed in Table S1 or by restriction digest of plasmids. mRNAs were transcribed using the mMESSAGE mMACHINE SP6 or T7 transcription Kit (Thermo Fisher Scientific). For preparing *tol2* mRNA, the *T7-TPase* plasmid (Addgene 51818) was used as a template. Injection mixes containing various concentrations of mRNAs were prepared with 0.5% (w/v) Phenol Red Solution prepared in PBS (Sigma-Aldrich, Darmstadt, Germany). Concentrations used for each mRNA were determined based on the previous report (Eimon et al., 2006) as below: *casp8* 150 ng/μL, *casp10* 150 ng/μL, *casp8l1* 150 ng/μL, *crmA* 250 ng/μL, *emiRFP670* 400 ng/μL, *tnfsf10l3* 100 ng/μL, *ripk1l* 150 ng/μL, *ripk3* 150 ng/μL and *ripk3-sfGFP* 150 ng/μL. 2-3 nL of the injection mix was injected in one-cell stage embryos. The quality of the mRNA was assessed right before injection through gel electrophoresis. To take images for the larval morphology, larvae were anesthetized with fish water containing 0.025% tricaine (Sigma-Aldrich) and mounted with 1.0% low-melting-temperature agarose (Thermo Scientific) prepared in fish water (0.18 g/L Instant Ocean Salt dissolved in RO water) on a 6-well glass-bottom dish (MatTek, MA, U.S.A., Cat# P06G-1.5-20-F). Larval bright-field images were taken using a Nikon Eclipse Ti-E inverted microscope equipped with a 4x objective lens (Nikon, Tokyo, Japan). Images were edited with ImageJ (National Institutes of Health). At around 3 hpf, dead and unfertilized embryos were removed from all experiments, and the fertilized healthy embryos were recorded as the initial number. The short-tail phenotype was recorded only for larvae with unambiguous reductions in tail length at 2 dpf as previously performed (Eimon et al., 2006). Embryonic death was recorded for embryos with completely dark appearance in the chorion at 1 dpf, and for larvae with no observed heartbeat at 5 dpf if the death was not evident.

### Generation of transgenic line

The *Tg(ef1a:crmA-P2A-EGFP)^cu78^* was generated by the Tol2 transposase-mediated transgenesis. Briefly, 25 ng/μL of *pTol2-ef1a-crmA-P2A-EGFP* plasmid was co-injected with 25 ng/μL of *tol2* mRNA into one-cell-stage embryos. Founder fish were identified by out-crossing with wild-type animals and observing F1 generation of larvae for EGFP expression. The lines from the founders that showed the strongest EGFP expression in the entire animal body were kept and used in this study.

### G0 CRISPR-Cas9 gene editing

CRISPR-Cas9-mediated G0 somatic mutagenesis was performed as previously described (Wu et al., 2018). 3 and 1 crRNAs were designed for targeting the exon 2 and 9 of *casp10*, respectively. 3 crRNAs were designed for targeting the exon 1 of *fadd.* All were designed using Custom Alt-R CRISPR-Cas9 guide RNA designing tool (Integrated DNA Technologies: IDT, IA, U.S.A.). Designed crRNAs, tracrRNA and Alt-R S.p. Cas9 Nuclease V3 were purchased from IDT. 1 crRNA targeting against *aicda* was used as a control for G0 CRISPR-Cas9 mutagenesis (Pagan et al., 2026). Since our G0 CRISPR-Cas9 experiments were conducted in the larval stages before the adaptive immune system is functional, *aicda* supposedly is not functional and thus served as a control gene to be targeted. crRNA targeting sequences are listed in Table S1. Cas9 RNP complex was prepared by following the manufacturer’s protocol. Briefly, in total 200 μM of crRNAs and 200 μM of tracrRNA were annealed in Nuclease Free Duplex Buffer (IDT). The final concentration of Cas9 protein and crRNA/tracrRNA in total in the injection mix were 0.83 μg/μL and 6 μM, respectively. Finally, Cas9 RNPs were activated at 37 °C for 10 min. 2-3 nL of activated Cas9 RNPs were immediately injected into one-cell stage embryos. To verify the gene-editing efficiency for *casp10*, and *fadd*, genomic DNA was extracted from randomly selected 10 Cas9 RNP-injected individual 5 dpf larvae using the HotShot method and High Resolution Melt (HRM)-based genotyping was conducted in a CFX Connect thermocycler (Bio-Rad) using the specific primers listed in Table S1. 100% of injected animals were verified to carry either *casp10* or *fadd* mutations observed by an obvious peak shift in the High Resolution Melt (HRM)-based genotyping assay (Fig. S2B, C).

### Protein sequence analysis

Protein sequence analysis was performed using the Ident and Sim software from The Sequence Manipulation Suite (Stothard, 2000). The access numbers for the analysed protein sequences were: *Homo sapiens* Casp8: Q14790 (NCBI); *Mus musculus* Casp8: ENSMUST00000191201.7 (Ensembl); *Danio rerio* Casp8: ENSDART00000104757.4 (Ensembl); *Danio rerio* Casp10: ENSDART00000186976.1 (Ensembl); *Danio rerio* Casp8l1: A0A2U9DQL0_DANRE (UniProt).

### Statistical analysis

Statistical analysis and visualization were performed on GraphPad Prism. All experiments were performed at least two times to ensure the reproducibility of the results, and one representative result is shown in the figures. The number of samples per group and the statistical test used are indicated in each figure legend or in the panels.

### Use of AI tools

AI tools were used to help drafting the manuscript and finding references in this study.

## Acknowledgements

We thank F.J. Roca for initially isolating the *casp8^cu76^* allele and for providing insights on the study, N. Goodwin and the University of Cambridge aquatic facility staff for zebrafish care, L. Whitworth and K. Shanahan for reviewing the manuscript, and the Ramakrishnan lab for insightful suggestions. Funding: This study was supported by a Wellcome Trust Principal Research Fellowship (223103/Z/21/Z) and NIH MERIT award R37 AI054503 (L.R.). N.Y. was supported from EMBO postdoctoral fellowship (ALTF1130-2021) and JSPS Overseas Research Fellowship (202460397). *pH2B-emiRFP670* was a gift from Vladislav Verkhusha (Addgene plasmid # 136571). *DR274* was a gift from Keith Joung (Addgene plasmid # 42250). *pSCAC-69* was a gift from Seok-Yong Choi (Addgene plasmid # 31241).

## Author Contribution Statement

J.F.M., L.R. and N.Y. contributed study concept and design, J.F.M. and N.Y. performed experiments and analysed data, J.F.M. and N.Y. generated resources for the study, L.R. and N.Y. supervised the study, J.F.M., L.R. and N.Y. wrote the original draft, J.F.M., L.R. and N.Y. edited the paper.

## Ethics Statement

This study was conducted in compliance with a guideline from the UK Home Office using protocols approved by the Animal Welfare and Ethical Review Body of University of Cambridge.

## Data Availability Statement

Zebrafish lines generated and plasmids used for it in this study can be obtained from the lead contact. Raw data reported in this work will be shared by the lead contact upon request.

