## Supplemental Information for "Zebrafish lack canonical necroptosis despite retaining functional RIPK3"

A

**Intercross of *casp8*$^{cu76}$ +/-**

| Genotype | Observed (#) | Observed (%) | Expected (%) |
|---|---|---|---|
| ***casp8* +/+** | 8 | 21.6 | 25 |
| ***casp8* +/-** | 19 | 51.4 | 50 |
| ***casp8* -/-** | 10 | 27.0 | 25 |

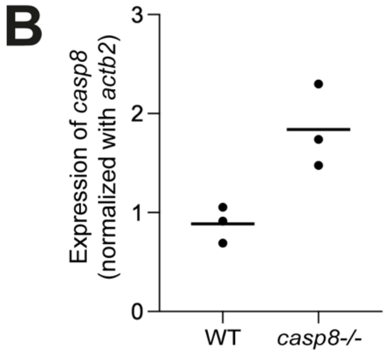

**Supplemental Figure 1**

(A) Observed and expected frequency of 14-week-old zebrafish with indicated *casp8*$^{cu76}$ genotype. (B) Lack of nonsense mediated decay of mRNA produced from *casp8*$^{cu76}$. The experiment was performed once with three biological replicates.

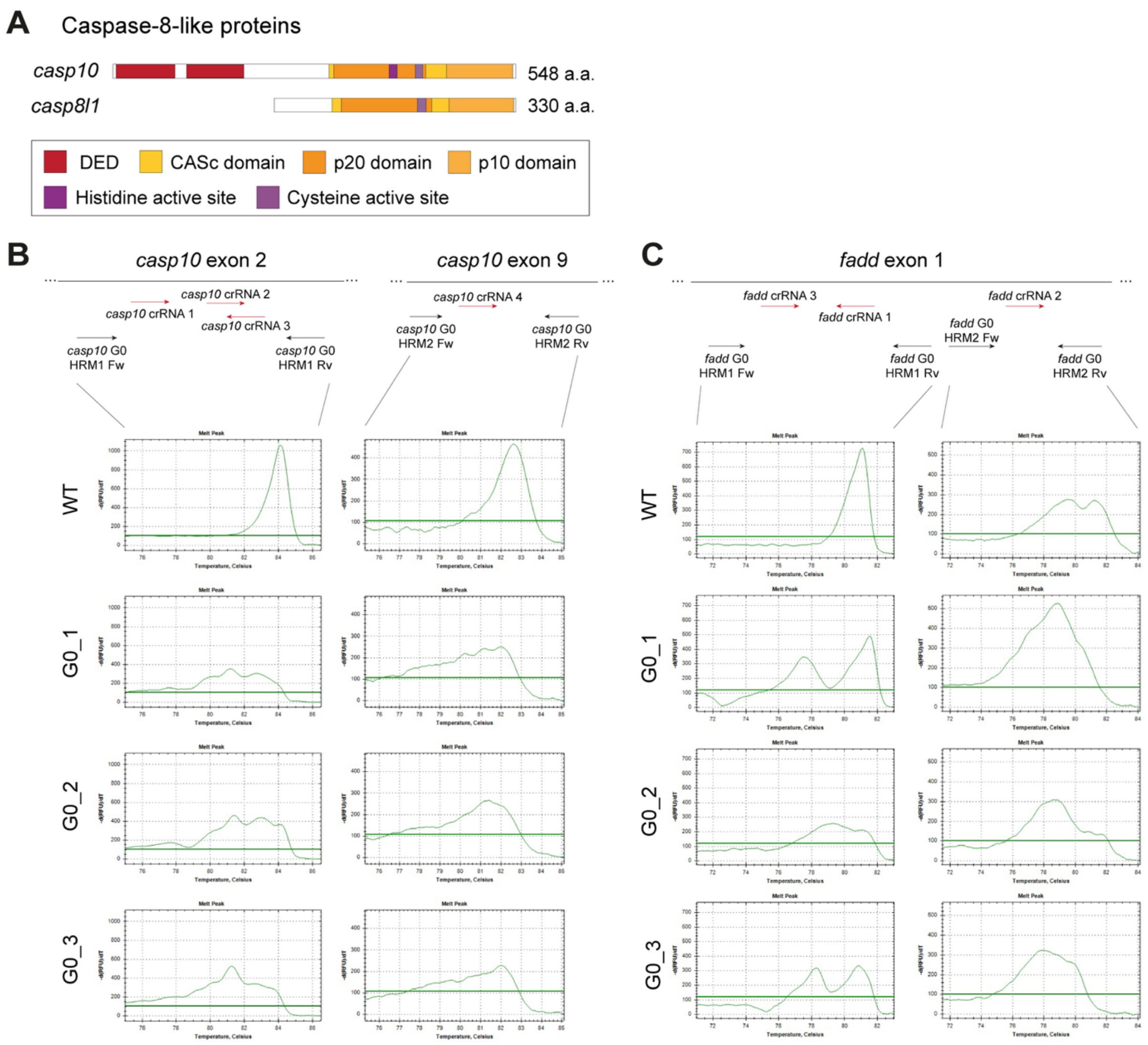

### Supplemental Figure 2

(A) Schematic representations of zebrafish Caspase-10 and Caspase-8l1 proteins. Example HRM genotyping patterns of *casp10* (B) and *fadd* (C) G0 CRISPR mutagenesis. WT: wild-type control. G0: Cas9 RNP injected animal.

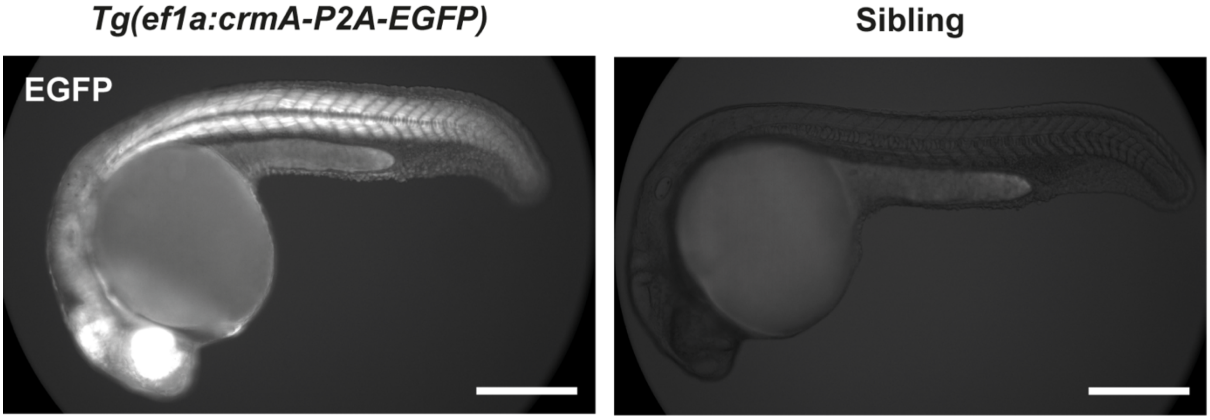

**Supplemental Figure 3**

Fluorescence images of a 1 dpf *Tg(ef1a:crmA-P2A-EGFP)*$^{cu78}$ embryo and its sibling.

Scale bar: 200 μm.

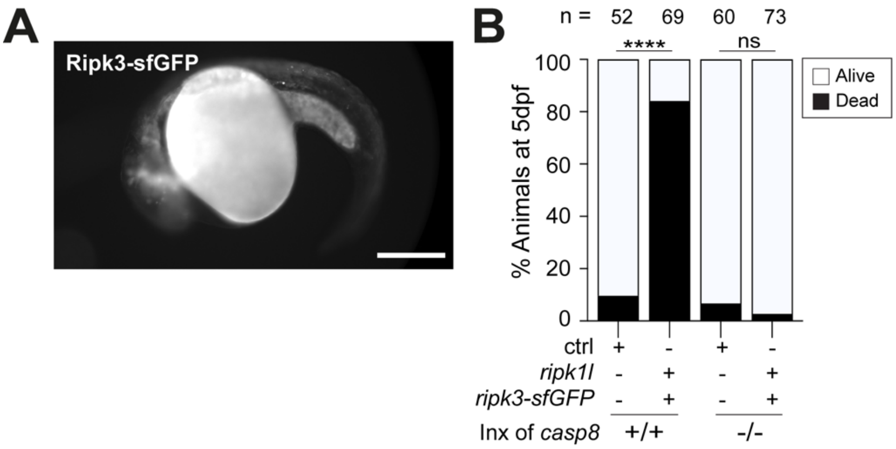

**Supplemental Figure 4**

(A) Fluorescent image of a *ripk3-sfGFP* mRNA injected 1 dpf embryo. Scale bar: 200 μm. (B) Percentage of larvae dead at 5 dpf from indicated crosses injected with combinations of *ripk1l*, *ripk3-sfGFP* and ctrl mRNAs. Inx: intercross. ctrl: *emiRFP670* mRNA. ***: *p*<0.0001 and ns: *p*>0.05 with Fisher's exact test.

### Supplemental Table 1

List of primers and crRNAs used in this study.

**Primers used in this study**

| Name | Sequence (5'-3') | Purpose |
|---|---|---|
| casp8-cu76_asp1 | GAAGGTGACCAAGTTCATGCTCTGGATGAGCAAGGATTGCT | To gentoype casp8^cu76 by KASP assay |
| casp8-cu76_asp2 | GAAGGTCGGAGTCAACGGATTCTGGATGAGCAAGGATTGAG | To gentoype casp8^cu76 by KASP assay |
| casp8-cu76_cp | AATGCGGCCAATTGTGAT | To gentoype casp8^cu76 by KASP assay |
| pCS2(+) open Fw | CAGACATGATAAGATACATTGATG | To open pCS2(+) plasmid for Gibson cloning |
| pCS2(+) open Rv | TGCAAAAAGAACAAGTAGCTTG | To open pCS2(+) plasmid for Gibson cloning |
| casp8 out Fw | CTTATACGCTACGAGGGTTC | Nested PCR to amplify casp8 CDS |
| casp8 out Rv | GTCCATCGTGTGTCCAGTC | Nested PCR to amplify casp8 CDS |
| casp8 Gibson Fw | agctacttgttctttttgcaATGGATCCTCAGATCTTTCAC | To clone pCS2(+)-casp8 |
| casp8 Gibson Rv | aatgtatcttatcatgtctgTCAGTCTATGGGCAGCACTAG | To clone pCS2(+)-casp8 |
| casp10 out Fw | GAACGCTTCACTGAAAGGAG | Nested PCR to amplify casp10 CDS |
| casp10 out Rv | AGTTTGCAGTTTGGAGCATC | Nested PCR to amplify casp10 CDS |
| casp10 Gibson Fw | agctacttgttctttttgcaATGGACGAGGAGATGGACATGGAC | To clone pCS2(+)-casp10 |
| casp10 Gibson Rv | aatgtatcttatcatgtctgTCAAGGAAGCCCTGGCGG | To clone pCS2(+)-casp10 |
| casp8l1 out Fw | AAGAGTGCATCAACACTTGG | Nested PCR to amplify casp8l1 CDS |
| casp8l1 out Rv | AACCATTGGCTTTGGTTCCT | Nested PCR to amplify casp8l1 CDS |
| casp8l1 Gibson Fw | acttgttctttttgcaATGGATAAAACTAGTAATCCTAACAAC | To clone pCS2(+)-casp8l1 |
| casp8l1 Gibson Rv | aatgtatcttatcatgtctgTTAGGAGACTCCATTCATGGG | To clone pCS2(+)-casp8l1 |
| ripk1l out Fw | GTGTGCTTTGAGAATAGAGG | Nested PCR to amplify ripk1l CDS |
| ripk1l out Rv | TGATAAAGACACAACAGTCG | Nested PCR to amplify ripk1l CDS |
| ripk1l Gibson Fw | agctacttgttctttttgcaATGTCTACTTCGGTCAGCCCGAGTCTC | To clone pCS2(+)-ripk1l |
| ripk1l Gibson Rv | aatgtatcttatcatgtctgTTACGGGGCGGCGGCGGAAT | To clone pCS2(+)-ripk1l |
| pCS2(+)-sfGFP open Fw | atggtgagcaagggcgagg | To open pCS2(+)-sfGFP plasmid for Gibson cloning |
| pCS2(+)-sfGFP open Rv | TGCAAAAAGAACAAGTAGCTTG | To open pCS2(+)-sfGFP plasmid for Gibson cloning |
| ripk3-sfGFP Gibson Fw | agctacttgttctttttgcaATGATCTGCTGTCATGAATTACAATC | To clone pCS2(+)-ripk3-sfGFP |
| ripk3-sfGFP Gibson Rv | CCTCGCCCTTGCTCACCATCCTACTGACATACGCAAAGCATTTC | To clone pCS2(+)-ripk3-sfGFP |

| | | |
|---|---|---|
| tnfsf10l3 out Fw | CTGTGAGTAGCGAGTCGAGC | Nested PCR to amplify tnfsf10l3 CDS |
| tnfsf10l3 out Rv | aagccccaaacacgtgtccc | Nested PCR to amplify tnfsf10l3 CDS |
| tnfsf10l3 Gibson Fw | agctacttgttctttttgcaATGGTGCAGCCTAAAAATCG | To clone pCS2(+)-tnfsf10l3 |
| tnfsf10l3 Gibson Rv | aatgtatcttatcatgtctgTCAGAGGTCAAACAGGAAGG | To clone pCS2(+)-tnfsf10l3 |
| T3 | attaaccctcactaaaggga | To linealize pCS2(+) plasmid by PCR for in vitro transcription |
| pCS2(+) linealize | gcggttagctccttcggtcc | To linealize pCS2(+) plasmid by PCR for in vitro transcription |
| casp10 G0 HRM1 Fw | GCTGTTTCTGTGCTCGGATCT | HRM genotyping for casp10 G0 CRISPR (guides AB, AA, AE; exon 2) |
| casp10 G0 HRM1 Rv | GTCCGTTTGGTGATGCTCAG | HRM genotyping for casp10 G0 CRISPR (guides AB, AA, AE; exon 2) |
| casp10 G0 HRM2 Fw | CAGTCTGCGGGACGTGT | HRM genotyping for casp10 G0 CRISPR (guide AH; exon 9) |
| casp10 G0 HRM2 Rv | AGCGCCTGCAGGATCTGA | HRM genotyping for casp10 G0 CRISPR (guide AH; exon 9) |
| faad G0 HRM1 Fw | GGATTCCGAGCAATGTTGC | HRM genotyping for fadd G0 CRISPR |
| faad G0 HRM1 Rv | CTCAGTGTTTTCAGGCCCAA | HRM genotyping for fadd G0 CRISPR |
| faad G0 HRM2 Fw | CGGTCAGACAGTTCTTATTGAAAA | HRM genotyping for fadd G0 CRISPR |
| faad G0 HRM2 Rv | ACCTGACTGACAGACATGAGAAT | HRM genotyping for fadd G0 CRISPR |
| aicda G0 HRM Fw | AATACGATATTTTGCTCCAGTGTGC | HRM genotyping for aicda G0 CRISPR |
| aicda G0 HRM Rv | CGATGCGTCGCTTTACTACAA | HRM genotyping for aicda G0 CRISPR |
| casp8 qPCR Fw | CTCAGCCTATAGAAAGATGC | RT-qPCR of casp8 |
| casp8 qPCR Rv | CAAGCCTCTGTTGTTTTTCC | RT-qPCR of casp8 |
| actb2 qPCR Fw | GTGCCCATCTACGAGGGTTA | RT-qPCR of actb2 |
| actb2 qPCR Rv | TCTCAGCTGTGGTGGTGAAG | RT-qPCR of actb2 |
| P2A-EGFP Fw | agcaggctggagacgtggaggagaaccctggacctatggtgagcaagggcgagg | To clone pTol2-ef1a-crmA-P2A-EGFP |
| EF1a Rv | ggtaagcttcagctagaactcgc | To clone pTol2-ef1a-crmA-P2A-EGFP |
| crmA Fw | agttctagctgaagcttaccATGGATATCTTCAGGGAAATCGC | To clone pTol2-ef1a-crmA-P2A-EGFP |
| crmA-P2A Rv | ctccacgtctccagcctgcttcagcaggctgaagttagtagctccgcttccggatccATTAGTTGTTGGAGAGCAATATCTACC | To clone pTol2-ef1a-crmA-P2A-EGFP |
| emiRFP670 Fw | atggcggaaggctccgtc | To clone pCS2(+)-emiRFP670 |
| emiRFP670-pA Rv | gtaatacgactcactatagtTTAgctctcaagcgcggtg | To clone pCS2(+)-emiRFP670 |

**crRNAs used in this study**

| Name | Target sequence (5'-3') | Target gene |
|---|---|---|
| casp10 crRNA 1 | GACCTCAACACTGTGAGCTC | casp10 |
| casp10 crRNA 2 | GAACCAGGACCTGCTGTCAT | casp10 |

| | | |
|---|---|---|
| casp10 crRNA 3 | ATGGATCCTCCAATGACAGC | casp10 |
| casp10 crRNA 4 | GCTTTGAGATCTTGACCCGT | casp10 |
| fadd crRNA 1 | GCACTCGAAGAGTTCAATCC | fadd |
| fadd crRNA 2 | ACGTCAAGAAGCCCGGCTGG | fadd |
| fadd crRNA 3 | CAAGCTTACAGATGATAACG | fadd |
| aicda | TCCACTATAAGAATGTGCGC | aicda |
